# PyCLM: programming-free, closed-loop microscopy for real-time measurement, segmentation, and optogenetic stimulation

**DOI:** 10.1101/2025.08.29.673155

**Authors:** Harrison R. Oatman, Beena C. Lad, Jared Toettcher

## Abstract

In cell biology, optical techniques are increasingly used to measure cells’ internal states (biosensors) and to stimulate cellular responses (optogenetics). Yet the design of all-optical experiments is often manual: a pre-determined stimulus pattern is applied to cells, biosensors are measured over time, and the resulting data is processed off-line. With the advent of machine learning for segmentation and tracking, it becomes possible to envision closed-loop experiments where real-time information about cells’ positions and states are used to dynamically determine optogenetic stimuli to alter or control their behavior. Here, we develop PyCLM, a Python-based suite of tools to enable real-time measurement, image segmentation, and optogenetic control of thousands of cells per experiment. PyCLM is designed to be as simple for the end user as possible, and multipoint experiments can be set up that combine a wide variety of imaging, image processing, and stimulation modalities without any programming. We showcase PyCLM on diverse applications: studying the effect of epidermal growth factor receptor activity waves on epithelial tissue movement, simultaneously stimulating ∼1,000 single cells to guide tissue flows, and performing real-time feedback control of cell-to-cell fluorescence heterogeneity. This tool will enable the next generation of dynamic experiments to probe cell and tissue properties, and provides a first step toward precise control of cell states at the tissue scale.

## Introduction

Optical techniques play increasingly important roles in modern cell biology. Imaging techniques have become increasingly sophisticated by pushing the limits of speed, resolution and 3D volumetric acquisition. Live-cell biosensors are available for a huge number of cellular processes from kinase activity to metabolism^1^, and optogenetic tools can now be used to activate protein binding, phase separation, cell signaling, and gene expression^2^. Rapid advances in artificial intelligence and machine learning for image processing have also made it possible to extract increasingly sophisticated information from biological imaging data^3^, opening the door to studying the behavior of large numbers of cells in complex geometries.

The rapid acceleration of probes, biosensors, and algorithms has driven a need to integrate all of these tools to enable real-time measurement and rule-based stimulation of large numbers of cells. Performing imaging and stimulation experiments on single cells or small collections of cells can be severely limiting in the study of dynamic, rare, or collective biological phenomena^4–6^. In these cases it rapidly becomes impractical or impossible to perform long-term stimulation experiments on one or a few cells at a time, to manually identify rare cells of interest and target them with individualized stimuli, or to simultaneously stimulate enough cells to initiate a collective tissue-scale response.

A growing number of “smart microscopy” approaches have been recently introduced to address these open challenges. Smart microscopy refers broadly to an emerging set of adaptive techniques in imaging and stimulation, empowering a microscope to autonomously decide how to image or stimulate a biological sample^7,8^. Kapitein and colleagues recently reported a system for performing real-time control over nucleocytoplasmic transport and movement of individual cells^9^. More recently, Pertz and colleagues extended image-based optogenetic stimulation across scales, from subcellular stimulation in single cells to multiple stimuli across populations of cells^10^. We reasoned that an ideal system should be reconfigurable for many different tasks, from single cell control to tissue-scale stimulation, and should be as simple to use as possible, without the need for any programming by the end user to combine any set of previously defined experimental workflows.

Here, we have developed a software package that accomplishes these goals called Closed-Loop Microscopy in Python (PyCLM). PyCLM combines fluorescent image collection through MicroManager^11^, cell segmentation through Cellpose^12^, and control of a digital micromirror device for optogenetic stimulation. Our system can be used to deliver programmatically-defined dynamic light inputs to multiple fields of view, track and stimulate large numbers of cells, and perform closed-loop experiments where real-time results of cell segmentation and biosensor activity are used to automatically update light inputs delivered to each cell. Any of these experimental modalities can be combined and configured independently for each field of view in a multi-point microscopy session using simple text-based configuration files and without end-user programming. We apply PyCLM to open questions in receptor tyrosine kinase (RTK)-driven cell migration. We use PyCLM to deliver traveling waves of light to cells expressing an optogenetic RTK, finding that cells exhibit optimal movement at intermediate wave speeds, and always migrate away from the source of waves. We also use PyCLM to deliver tailored subcellular light stimuli to each cell within a tissue, finding that comparable migratory responses can be achieved by both single-cell stimuli and tissue-scale waves. Finally, we showcase how PyCLM can implement closed-loop feedback control on fluorescence intensity to counteract cell-to-cell variability. We anticipate that this toolbox will be broadly useful for probing dynamic single-cell biology at large scales and controlling cell behavior to drive desired tissue-scale outcomes.

## Results

### The PyCLM architecture for closed-loop imaging and optogenetic stimulation

Our overall goal was to develop software to enable complex imaging and stimulation experiments for cell and developmental biology (**Fig. 1A**). Current optogenetic experiments often require the ability to deliver dynamic light patterns, such as traveling waves or light gradients for the study of tissue migration or developmental patterning^13,14^, yet specification of these dynamic patterns remains difficult in standard software packages. We also sought to integrate real-time image segmentation and light stimulation, so that the location and intensity of stimulation could be tailored to the geometry of specific samples (e.g. embryos or tissues). This “closing the loop” would allow light stimulation to adaptively respond in a programmatically predetermined way as an experiment progresses.

**Fig. 1:**
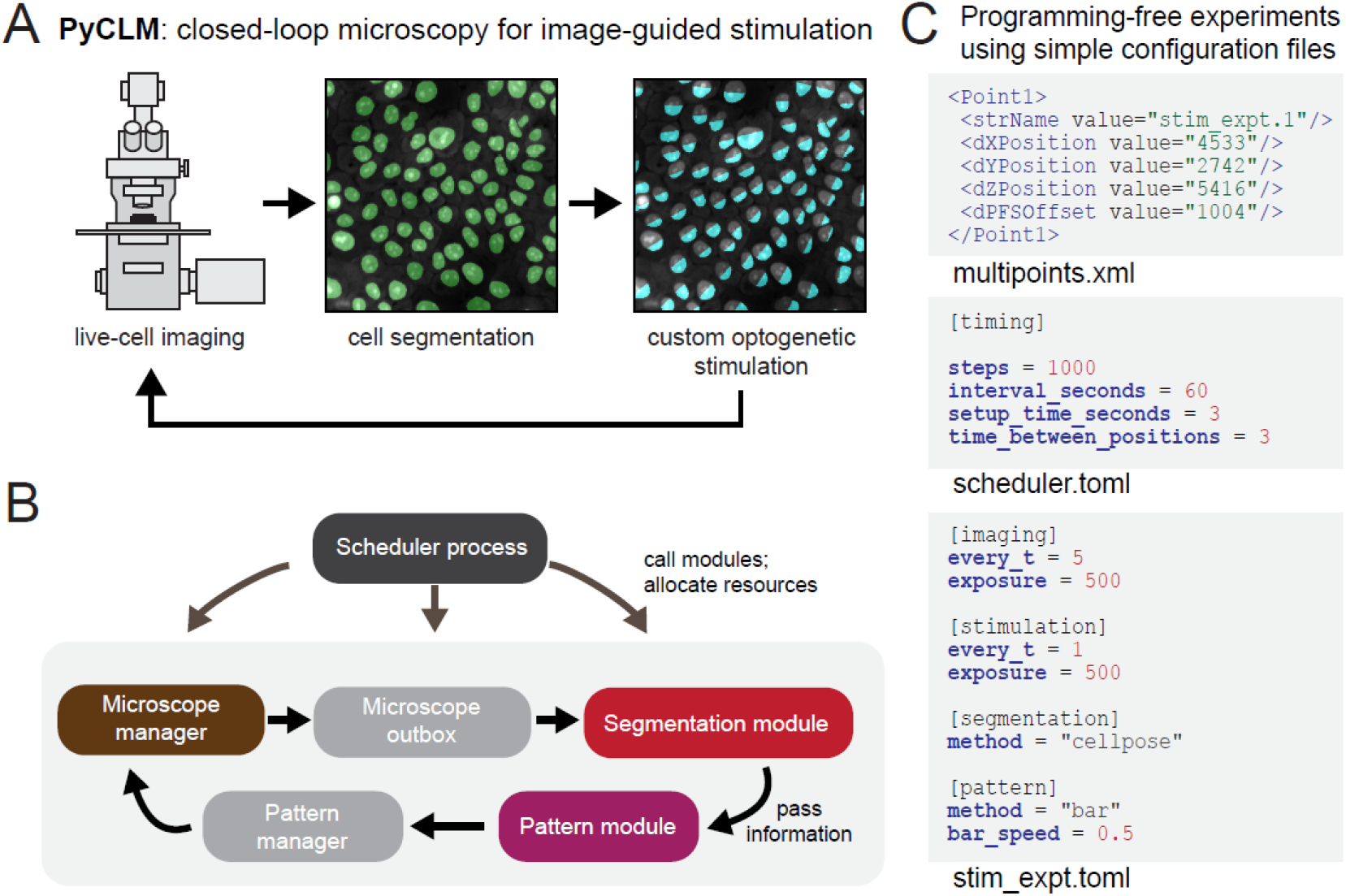
Overview of the PyCLM closed-loop microscopy pipeline. (**A**) PyCLM is designed to combine imaging, cell segmentation, and delivery of customized optogenetic patterns to many cells across multiple fields of view. (**B**) Organization of the PyCLM software. A master scheduler controls the sequencing of all modules. Five modules pass information in a feedback loop: a microscope manager for hardware control, an “outbox” for image data storage, a segmentation module for identifying biological structures, a pattern module for implementing rules for optogenetic stimulation, and a pattern manager that collects patterns and delivers them back to the hardware manager. Multiple microscope manager, segmentation, and pattern modules can be combined in a single experiment to run on different fields of view, and can be easily interchanged to implement different experimental designs (e.g. segmentation algorithms or stimulus designs including dynamic light patterns or feedback-controlled illumination rules). (**C**) A example of the configuration files required to run an experiment. A multipoints.xml file provides the names and spatial coordinates of each imaging position. Each position’s name is linked to a.TOML configuration file that defines a combination of modules and their configurations so that multiple complex experiments can be run in parallel. The scheduler.toml configuration file specifies overall settings for the entire experiment.

In achieving this overall goal of closed-loop microscopy, we were guided by several practical design considerations. First, we sought to achieve a high degree of flexibility, so that many different experimental workflows could be performed on different fields of view in a single imaging session. Next, we aimed to build an extensible system, so that diverse experiments could be designed and implemented on top of the existing software. For example, one must be able to swap in and out different cell segmentation models, apply different time-varying light input patterns to each field of view, and implement a broad range of possible feedback strategies. Finally, we sought the simplest possible interface, so that users without programming background could remix any existing combination of modules to define new experiments. To produce such an accessible and generalizable workflow, we brought together excellent resources that are already used in a diverse set of experimental systems and image processing tasks: Cellpose for generalist image segmentation, and MicroManager for microscope control and optogenetic stimulation.

One challenge in our design is scheduling: to satisfy the closed-loop design, cell segmentation and optogenetic stimuli must be performed before the next round of imaging. Our solution to this challenge is an architecture where processes (imaging, cell segmentation, stimulation) are implemented as distinct modules that run as independent threads (**Fig. 1B**). A master scheduler process manages each of the modules to ensure appropriate sequencing of events. We implemented a scheduler and five classes of modules: (1) the “microscope manager” that works with MicroManager to control stage movement, image acquisition and optogenetic stimulation, (2) the “microscope outbox” that controls data storage and handles images, (3) “segmentation module” that implements image segmentation, (4) the “pattern module” that defines any dynamic illumination patterns or updating rules (e.g. feedback control), and (5) the “pattern manager” that stores patterns and provides them to the microscope manager as needed. The user may configure or swap three of these modules – the microscope manager, segmentation module, and pattern module – as needed to implement different experimental designs. Crucially, each field of view uses its own specific combination of modules and configuration parameters, so that many different experiments can be run in parallel during a single multi-point acquisition.

From the end-user perspective, running a PyCLM workflow is performed not by writing Python code, but by editing a set of human-readable Tom’s Obvious, Minimal Language (TOML) configuration files. These.TOML files allow the user to set any parameters needed for imaging, segmentation, and feedback control. Each multipoint field of view is associated with its own.TOML file that describes the specific experiment to be performed at that position, and a master `schedule.toml` file sets the overall experiment length and imaging frequency. The user thus needs to provide a set of imaging coordinates, a TOML configuration file for each position, and a scheduler configuration file to specify a complete experiment (see examples of this code in **Fig. 1C**). We provide.TOMLs for running all experiments described in this manuscript as supplementary files.

### Dynamic control of optogenetic stimuli enables directional tissue movement

As a first application of PyCLM, we set out to deliver traveling waves of optogenetic stimulation to a tissue to study how dynamic patterns of receptor tyrosine kinase (RTK) activity affect tissue movement. Recent studies have revealed that developmental and regenerative processes in epithelial tissues are often accompanied by propagating waves of receptor tyrosine kinase (RTK) signaling (**Fig. 2A**). Fibroblast growth factor (FGFR) signaling waves travel across regenerating zebrafish scales^15^, and waves of epidermal growth factor receptor (EGFR) activity are prominent in many mammalian epithelial tissues^16^. The reported wave speed varies over a broad range between tissue contexts (zebrafish scale regeneration: 2-10 μm/h; mouse epidermal tissue: 120 μm/h)^15–17^. It has been hypothesized that these traveling waves of RTK signaling serve to organize collective cell movement toward sites of tissue damage. Multiple prior optogenetic studies have revealed that cell movement can be driven downstream of local RTK pathway stimulation^14,18,19^. However, in some cases cells move “upstream” toward the source of the activity wave, whereas in other cases the direction is reversed, with cell movement following the activity wave. A systematic study of how RTK activity waves at different speeds organize tissue movement has not been performed.

**Fig. 2:**
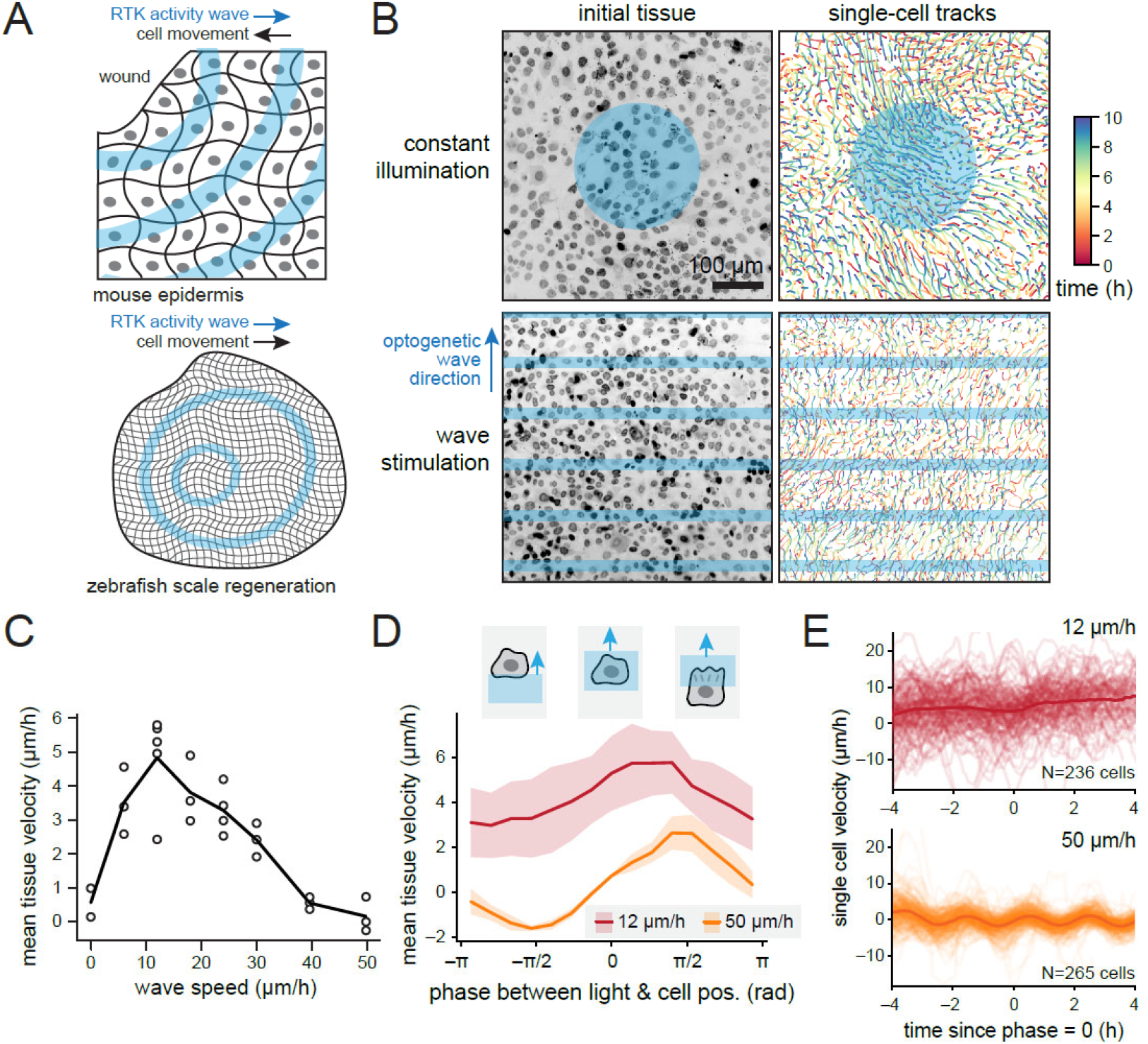
Waves of OptoEGFR stimulation drive rapid, directed tissue flows. (**A**) Tissue-scale waves of receptor tyrosine kinase activity have been observed in many systems, where they are thought to control cell movement. However, wave speed and movement direction are both highly variable between cellular contexts, with migration toward the source of waves in the mouse epidermal wound healing and away from the source in zebrafish scale regeneration. (**B**) In cells expressing an optogenetic EGF receptor (OptoEGFR), constant illumination drives inward tissue movement as previously observed. Waves of illumination drive robust tissue movement away from wave source. (**C**) Quantification of tissue velocity as a function of wave speed across independent replicates. (**D**) Quantification of tissue velocity as a function of the phase of the light wave relative to each tissue position. Data shows mean + S.D. from n=5,3 replicates in the 12 and 50 μm/h wave speeds, respectively. (**E**) Quantification of single-cell velocities as a function of time since complete overlap by the light bar (defined as phase = 0) for 12 and 50 μm/h light waves. Data shows single cell traces and their mean for n=236,265 analyzed and tracked cells, respectively. See also Fig. S1.

To enable light-based control of RTK activity, we used a previously described light-activated EGF receptor (OptoEGFR)^18^ which is based on receptor clustering using the 450 nm blue light sensitive OptoDroplet system^20^. We generated a MCF10A OptoEGFR-expressing cell line^21^ and incubated these cells with an infrared-fluorescent Hoechst dye for tracking their nuclei during imaging experiments. We first confirmed that these OptoEGFR cells migrated in response to constant light stimuli as observed previously^18^ (**Fig. 2B**, top panels; **Figure S1A**; **Movie S1**). We then wrote a PyCLM module to stimulate cells with traveling horizontal “bars” of light. The `bar` module implements these dynamic patterns with simple parameters that can be set in the `bar.toml` configuration file: the spatial period between successive bars, the wave speed, and the duty cycle (or proportion of the period that is illuminated versus dark). Blue light patterns were applied once per minute to maintain optogenetic activation, whereas fluorescent images were collected once every 5 min to monitor cell positions. The flexibility of our interface enabled us to stimulate up to 15 different fields of view with differently-parameterized light waves in a single experiment. We sampled a broad range of wave speeds from 8-50 μm/h and performed each experiment for 16 h.

We found that traveling waves of illumination indeed produced robust tissue movement (**Fig. 2B**, bottom panels). We always observed tissue movement in the direction of the traveling wave, consistent with *in vivo* observations in regenerating zebrafish scales^15^ but contrary to the migration direction observed in wounded mouse epidermis^16^. Quantifying the results across different optogenetic wave speeds revealed a maximum tissue velocity of ∼5 μm/h at a wave speed of 12 μm/h, with decreasing tissue movement observed as the RTK activity wave speed was increased (**Fig. 2C**; **Movie S2**). No overall translational movement of the tissue was observed for fast (50 μm/h) wave speeds (**Movie S3**).

How does the movement of a cell vary as a wave of RTK activity sweeps over it? Addressing this question is challenging because the traveling waves are centered at different positions in the tissue at each time point, and cells may also move. To account for these effects, we computed the phase of the traveling wave at each time and spatial position, assigning a phase of zero when the light wave is centered on the position. We then plotted the velocity of all cells as a function of the phase of the light input that they experience (**Fig. 2D**; **Fig. S2B-C**). We found that for fast (50 μm/h) waves, cell movement was symmetric about zero, with cells moving toward an incoming light wave as it approaches, and following the outgoing wave as it recedes. In contrast, for slow (12 μm/h) waves, cells consistently moved in the outgoing wave direction at all phases, but moved most rapidly at phases when partially illuminated by the outgoing wave. This data suggests that for slow wave speeds, partially illuminated cells are able to polarize and move in the direction of the light wave, directing the motion of the entire tissue, whereas for fast wave speeds, cells oscillate in their polarization direction with each wave without achieving sustained movement. Similar results were also observed from plots of individual cell movement over time, and centered at the time where each cell was at zero phase relative to the light input (**Fig. 2E**). Here, the sinusoidal motion of cells with each incoming light wave was apparent at 50 μm/h but not 12 μm/h, likely because many single cells migrate at a similar speed to the outgoing wave and thus only change relative phase slowly over time.

Taken together, PyCLM enabled the easy design of experiments that implemented traveling waves of optogenetic stimulation to tissues. These experiments revealed that traveling waves of RTK activity can indeed drive efficient tissue movement. Our data reveal that RTK activity waves are particularly effective at slow wave speeds, and predominantly lead to cell movement in the direction of the outgoing wave. These data suggest that other factors are likely to be required to explain cell migration “upstream” toward wounds in the presence of rapid RTK activity waves such as those observed in the mouse epidermis. These factors may include differences in cell density or tissue mechanics near wounds, or other signaling cues produced at sites of injury.

### Closed-loop segmentation and stimulation enables control over complex tissue flows

In the preceding section we examined tissue movement in response to “supracellular” optogenetic stimuli: waves of stimulation whose spatial scales are larger than that of individual cells. This approach differs from the classic experiment for optogenetic cell migration, where a light input is tailored to one side of a cell to drive its directed motion^19,22–25^. How would features of migration differ between these modes? While subcellular stimuli have been widely used to control cell migration, their use is made cumbersome by the requirement to manually place an illumination pattern on a cell that must be updated as it moves. Scaling this approach up to hundreds or thousands of moving cells would quickly become untenable, and could represent an ideal use case for an automated microscopy approach.

To enable local stimulation of all cells in a field of view, we implemented a segmentation module that uses Cellpose’s CPSAM model^26^ to identify the position of all nuclei using their Hoechst nuclear dye signal. We then implemented a stimulation module that uses the segmentation masks to construct a Voronoi tessellation to identify regions associated with all cells. We implemented a stimulation rule to apply a gradient of light oriented in a counter-clockwise direction relative to the center of the field of view, based on our hypothesis that these inputs would drive a counterclockwise rotational flow of the entire tissue (**Fig. 3A**). Rotational fluid or tissue flows are found in various biological contexts including left-right patterning^27^ and hair follicle formation^28^, yet directly programming rotational motion in cellular systems has not yet been achieved.

**Fig. 3:**
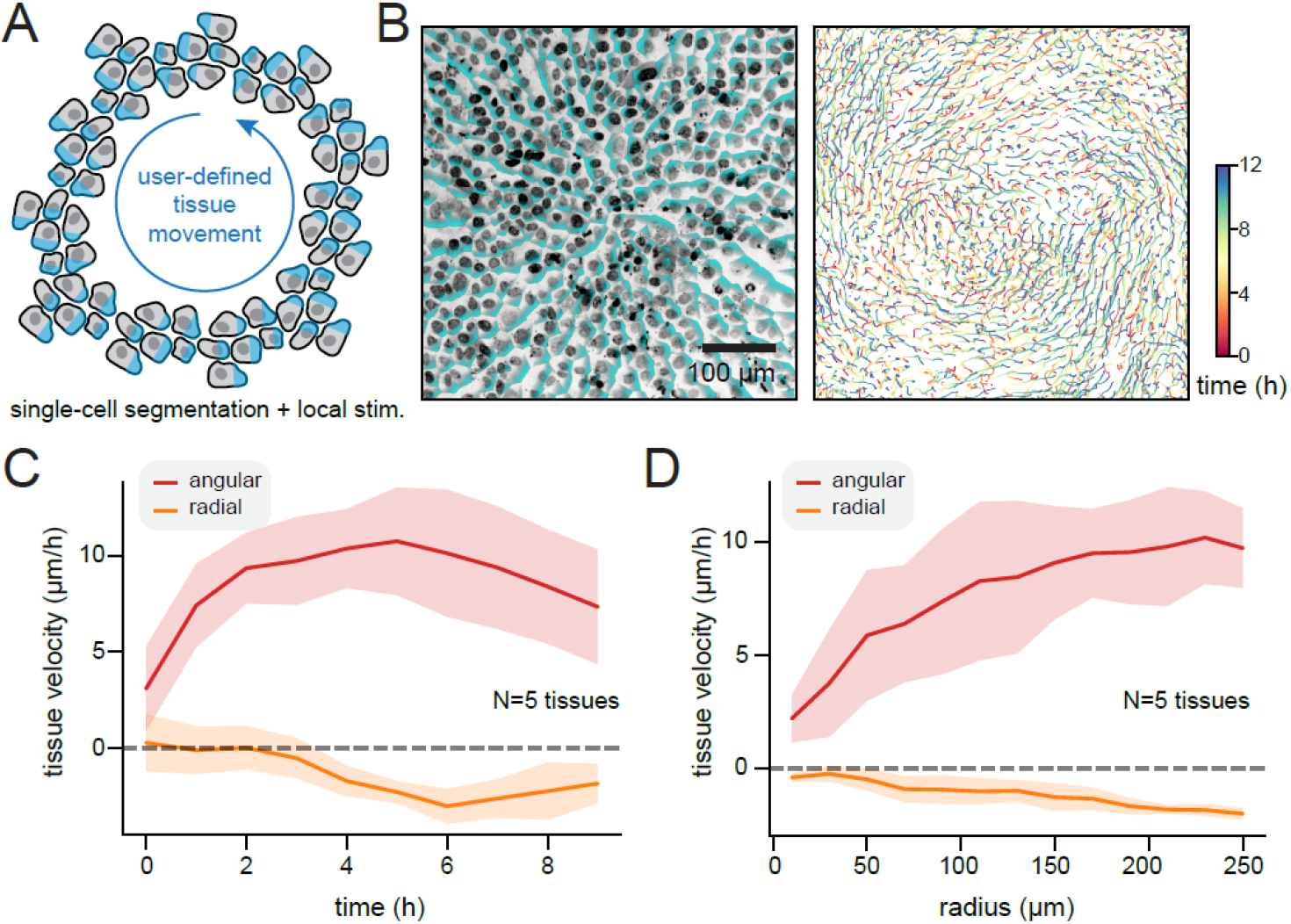
Closed-loop directional stimulation of large cell populations produces user-directed tissue flows. (**A**) As an alternative way to control tissue movement, we envisioned an experiment where all individual cells in an epithelial tissue are locally illuminated to drive their polarization and outgrowth. If such stimuli are applied tangent to cell’s position from an origin, the tissue would be expected to move in a rotational flow. (**B**) Left: Image of live MCF10A tissue stained with a Hoechst nuclear dye, with the illumination pattern overlaid in teal. Right: Cell movement tracks over time for the experiment described in **A**. (**C**) Quantification of cell velocities as a function of time between n=5 tissue replicates, separated into angular and radial components. Movement is rapid in the angular direction, with a slight inward motion in the radial direction. (**D**) Quantification of cell velocities as a function of radial distance from the tissue origin between n=5 tissue replicates, showing faster cell movement at positions far from the origin.

Indeed, we found that our `vortex` module drove robust rotational motion of an MCF10A OptoEGFR-expressing epithelial monolayer (**Fig. 3B**; **Movie S4**). We were able to segment and design local stimuli to more than eight fields of view within 5 mins, the time between successive rounds of imaging, while simultaneously maintaining a light stimulation frequency of 1/min. Quantifying cell movement over time revealed robust counterclockwise motion, with a small amount of inward migration indicated by a negative radial velocity (**Fig. 3C**). We also observed that the angular velocity of tissue movement decreased at interior tissue positions (**Fig. 3D**).

Interestingly, the typical migration speeds achieved by stimulating each cell with a customized light gradient ranged between 5-10 μm/h, comparable to what was achieved by supracellular traveling waves (**Fig. 2C**), and suggesting that waves achieve approximately maximal tissue movement speeds. Overall, these data demonstrate that PyCLM can be applied to study a light-controlled cell migration in a variety of contexts. Our experiments reveal that receptor tyrosine kinases can achieve similar tissue-scale migration outcomes through supracellular dynamic waves or local single-cell gradients.

### PyCLM can be used for real-time feedback control of cell-to-cell heterogeneity

Cell-to-cell heterogeneity is ubiquitous in cell biology, affecting cells’ response to signaling cues^29^, gene expression^30^, and phenotypic properties such as chemotherapeutic drug resistance^5^. Optogenetic feedback control can in principle mitigate some of this heterogeneity by delivering tailored inputs to each cell to compensate for intrinsic differences in gene expression or cell state^31,32^. However, a challenge arises at scaling up this kind of control. For large populations of cells, each cell must be segmented, its state measured, and an appropriate light input must be computed that drives its state toward a desired value.

We reasoned that PyCLM’s capabilities also make it ideal for implementing feedback control on large populations of single cells. To accomplish this goal, we combined our CPSAM segmentation module with a new module that implements “bang-bang” feedback control over each nucleus’ fluorescence intensity. Bang-bang control is a feedback control strategy well known from residential thermostats (e.g., heat is applied whenever the room temperature is less than a desired value, and off when it is greater than value).

As a test case for biological control, we cultured MCF10A cells that expressed a TagRFP-H2B nuclear marker at variable expression levels. This system provides an ideal case for blue light-dependent control due to the fact that like many RFPs, TagRFP is photochromic, exhibiting a change in red fluorescence after blue light exposure^33,34^. Indeed, performing time-lapse imaging of TagRFP-H2B expressing MCF10A cells revealed that their nuclear fluorescence increased by approximately 4.5-fold following illumination by 450 nm light. This change is reversible upon removal of the blue light stimulus (**Fig. 4B**; **Movie S5**). We reasoned that blue light stimuli could be tailored to each cell to overcome the initial cell-to-cell heterogeneity in TagRFP expression, producing tightly controlled cellular fluorescence across the field.

**Fig. 4:**
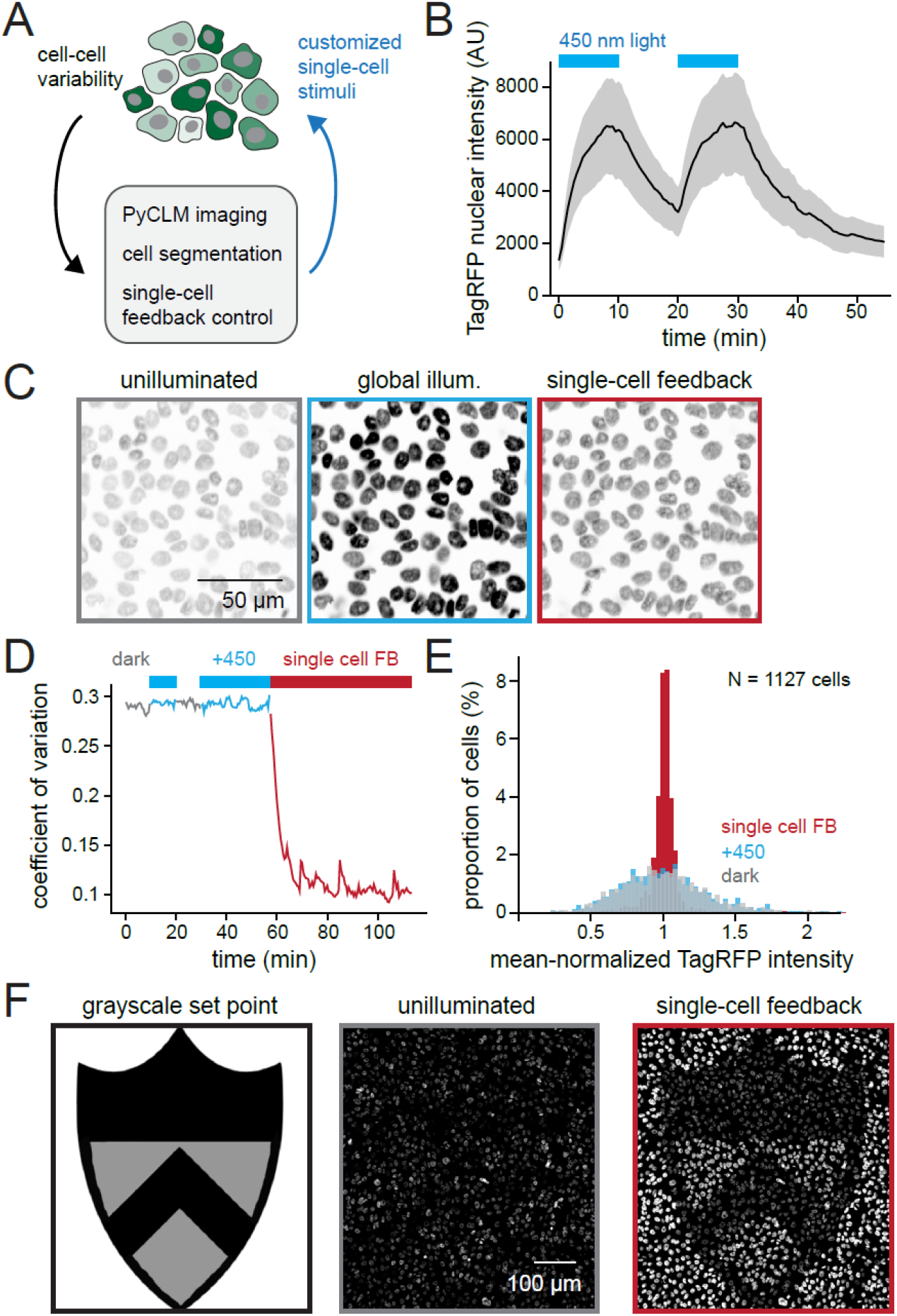
A “bang-bang” optogenetic feedback controller for cell populations. (**A**) Because it integrates real-time analysis and optogenetic stimulation, PyCLM can be used for real-time feedback control of cellular properties. We envisioned an experiment where heterogeneous fluorescent signals could be “normalized” by tailoring light inputs delivered to each cell. (**B**) MCF10A cells expressing a nuclear TagRFP-NLS construct were exposed to blue light or darkness and their red fluorescence was measured over time. A four-fold changes in TagRFP fluorescence in the presence or absence of blue light due to photochromicity of this fluorescent protein. Curves show mean + S.D. from n=1127 cells. (**C**) Images from a field of TagRFP-expressing cells in the dark, under global blue light, or subjected to a bang-bang feedback controller that applies a blue light stimulus to each cell if its intensity falls below a set point. (D) The coefficient of variation in TagRFP fluorescence calculated between n=1127 cells in the field of view is plotted as a function of time under conditions of darkness, global blue light, or feedback control. (E) Histogram of nuclear TagRFP fluorescence intensities in n=1127 cells in dark, light and under feedback control. (F) Feedback control to produce a spatial of TagRFP fluorescence based on a grayscale image of set points. Left: the set point image. Middle: an un-illuminated field of MCF10A cells. Right: the same cells after feedback control is applied. See also Fig. S2.

We combined PyCLM modules to image MCF10A TagRFP-H2B cells every 30 sec, segment each nucleus, and apply feedback-controlled inputs to drive the red fluorescence of all cells in the field of view to a uniform set point (**Fig. 4C**; **Movie S6**). Global illumination with a uniform blue light input increased the brightness of all cells (**Fig. 4C**, left vs middle) but cells retained a high degree of variability in fluorescence. In contrast, local feedback control was able to compensate for cells’ intrinsic differences in fluorescence (**Fig. 4C**, right), driving uniform nuclear fluorescence in all cells. We quantified cell-to-cell variability by the coefficient of variation (CV) of cells’ nuclear TagRFP intensity under different global conditions (dark vs blue light) and in the presence of feedback control (**Fig. 4D**) as well as by examining the overall distribution of single-cell intensities (**Fig. 4E**). Global illumination had no effect on the CV, whereas feedback control drove a three-fold reduction in CV across the cell population within 20 min that was maintained for the duration of the control experiment (**Fig. 4D**), an effect that was also apparent from the tight distribution of single-cell intensities achieved by feedback control (**Fig. 4E**; **Fig. S2A**). Despite all cells attaining the same final set point, we found that the amount of blue illumination delivered to each cell by the controller scaled inversely with the cell’s initial fluorescence (**Fig. S2B-C**), indicating that the information about cell-to-cell heterogeneity was now stored within the controller inputs, rather than fluorescent output.

Our PyCLM implementation of feedback control is quite general, and also enables set points to be chosen locally at different positions in a field of view. We showcased this functionality by providing a grayscale set point image as input, driving the nuclear fluorescence intensity of a field of cells to match the desired intensity pattern (**Fig. 4F**; **Movie S7**). Taken together, these data demonstrate that PyCLM provides a simple, extensible interface to implement intensity-based optogenetic feedback control on large numbers of cells for reducing cell-to-cell variability or controlling cell states.

## Discussion

Smart microscopy” is gaining prominence as a transformational technology for cell biology studies. The smart microscopy revolution is driven by numerous recent advances. Artificial intelligence and machine learning make it possible to automate many computational tasks that previously required extensive human intervention, opening the door to real-time cell segmentation, tracking, and analysis. At the same time, an increasing library of biosensors makes it possible to measure ever more traits of the cells being studied. Finally, optogenetic tools provide optical actuators, enabling experimentalists to take advantage of the information gained by imaging to design new perturbations in real-time. Here, we present a software environment for combining each of these modalities to perform dynamic stimulation experiments and automate detection and perturbation experiments on large numbers of cells.

Many of the examples we draw upon in this study come from optogenetic control of cell migration. Light-controlled single cell migration is one of the classic applications of optogenetics in cell biology, and indeed was demonstrated in some of the earliest papers in the field^22,24^. However, these tools have only recently been applied to large-scale tissue movement and collective cell migration^14,18^, and major questions remain about how mechanical and chemical information is propagated across tissues. Here we show that PyCLM can perform important classes of experiments for optogenetic studies of collective cell migration: sweeping light inputs across a tissue to recapitulate dynamics observed *in vivo* and locally stimulating single cells at large scales to reveal how the collective polarization of many single cells can produce large-scale tissue flows.

We find that waves of light-activated EGFR activity are indeed sufficient to drive tissue movement that is oriented along the outgoing wave at speeds similar to what is achieved by locally stimulating every cell in the tissue. These migration data are consistent with what is observed in zebrafish scale regeneration: directed movement along RTK activity waves of 2-10 μm/h^15^. In contrast, studies from mouse epidermal regeneration and cultured Madin-Darby canine kidney (MDCK) cells reported rapid (>100 μm/h) waves of EGFR activity that are oriented in the opposite direction of tissue expansion^14,16^. Our data would suggest that EGFR activity waves are unlikely to be solely responsible for directed cell migration in these contexts. Indeed, previous studies show that MDCK tissues still undergo migration toward unoccupied regions in the presence of EGFR pathway inhibitors that extinguish the activity waves originating from these regions^14,17^, suggesting that other directional cues also play a role in biasing overall tissue motion.

We further demonstrate that PyCLM is able to perform closed-loop optogenetic stimulation by acquiring images, segmenting biological features of interest, and using these features to compute new light stimuli in real time during experiments. We apply this capability to locally stimulating cell migration and performing feedback control on single-cell fluorescence intensity across populations of more than 1,000 cells. However, one limitation of our current implementation is that it does not track and store information about single cells’ behavior over time. Tracking single cells over time would be useful for certain calculations, such as identifying cells with different behavior over time (e.g. fast versus slow migrators) or implementing feedback control algorithms that require knowledge of each agent’s history (e.g., integral feedback controllers that integrate accumulated error over time). We aim to add support for cell tracking in future versions of PyCLM, as well as control of multiple light sources to perform still more complex multi-input, multi-output control experiments.

Smart microscopy aims to endow imaging systems with the ability to perform complex experiments without extensive real-time human intervention. Our PyCLM package provides a simple interface for performing one class of smart microscopy experiments: imaging, segmentation, and optogenetic stimulation. Our tool, along with others in this emerging field, will likely enable the next generation of complex imaging-based experiments for biological discovery and control.

## Supporting information

Movie S1 - circle stimulation

Movie S2 - 12 um/h wave stimulation

Movie S3 - 50 um/h wave stimulation

Movie S4 - "vortex" stimulation

Movie S5 - TagRFP blue light toggle

Movie S6 - TagRFP feedback control

Movie S7 - TagRFP spatial feedback patterning

## Acknowledgements

We thank Joachim Goedhart for valuable discussions about photochromicity of red fluorescent proteins, and all members of the Toettcher lab for helpful comments throughout the project. This work was supported by NIH grant R01GM144362 and NSF RECODE grant 2134935 (to J.E.T.), NIH training grant T32GM148739 and an NSF GRFP fellowship DGE-2444107 (to B.C.L.), and NIH training grant T32HG003284 (to H.R.O.).

## Author contributions

H.R.O. and J.E.T. conceptualized the project. H.R.O. wrote the code and performed all data analysis. B.L. and J.E.T. performed all experiments. H.R.O. and J.E.T. wrote the paper with editing from all authors. J.E.T. supervised the research.

## Declaration of interests

J.E.T. is a scientific advisor for Prolific Machines and Nereid Therapeutics. The remaining authors declare no conflicts of interest.

## Materials and Methods

### Software

PyCLM was written in Python v3.11.5 using pymmcore-plus v0.13.7 and Cellpose^12^ v4.0.6 for microscope hardware control and cell segmentation, respectively. MicroManager Studio Version 2.0.3 was configured for the microscopy setup described below. All data analysis was performed using custom scripts written in Python. All code and software requirements are available on the project’s Github page (www.github.com/Harrison-Oatman/PyCLM). Pymmcore-plus is available at https://github.com/pymmcore-plus.

### Tissue culture

MCF10A-5E cells^35^ were cultured in DMEM/F12 media supplemented with 5% horse serum (Invitrogen, 16050122), 20 ng/mL EGF (Peprotech, AF-100-15-1MG), 0.5 µg/mL hydrocortisone (Sigma-Aldrich, H0888), 100 ng/mL cholera toxin (Sigma-Aldrich, C8052), 10 µg/mL insulin (Sigma-Aldrich), and 1% penicillin/streptomycin. All cells were maintained at 37°C and 5% CO2. HEK293T cells were cultured in DMEM supplemented with 10% fetal bovine serum (R&D Systems, 26140079) and 1% penicillin/streptomycin (Gibco,15140122).

### Cell preparation for microscopy

Cells were imaged on glass-bottomed, black-walled 96-well plates (Cellvis, P96-1.5H-N) coated with fibronectin. Wells of 96-well plates were first incubated with 10 µg/mL fibronectin dissolved in PBS at 37°C for a minimum of 30 min. After fibronectin coating, wells were washed twice with DI water and seeded at 30,000 cells/well 1 day prior to imaging. For improved adhesion, cells were plated into 150 μL of media, then spun down in a tabletop centrifuge for 30 sec. 3 hours prior to imaging, growth medium was replaced with low-growth factor “starvation” medium. Starvation medium was identical to growth medium, but lacked horse serum, EGF, and insulin, and was supplemented with 3 mg/mL bovine serum albumin. At least 1 hour prior to imaging, medium was replaced with starvation medium containing 0.1uM nuclear dye Hoechst Janelia fluor 646 (Tocris, 6804). All cells were maintained at 37°C and 5% CO2.

### Imaging

All microscopy experiments were performed on a Nikon Eclipse Ti-2 microscope using a Crest V3 spinning disk confocal system, Celesta 545 and 638 nm lasers, a Kinetix 10MP CMOS camera, and a Mightex Polygon 1000 DMD driven by a Sola LED light source. For Fig 2 and 3, 450 nm light stimuli were applied every 1 min at 10 mW/cm^2^ for 500 msec; for Fig 4, 450 nm light stimuli were applied every 30 sec at 30 mW/cm^2^ for 500 msec. Images were collected using a 20X air objective lens.

## Notes

https://github.com/Harrison-Oatman/PyCLM

