## Supplementary Information for "PyCLM: programming-free, closed-loop microscopy for real-time measurement, segmentation, and optogenetic stimulation"

### Contents

Supplementary Figures 1-2

Supplementary Text

Supplementary Movie Legends

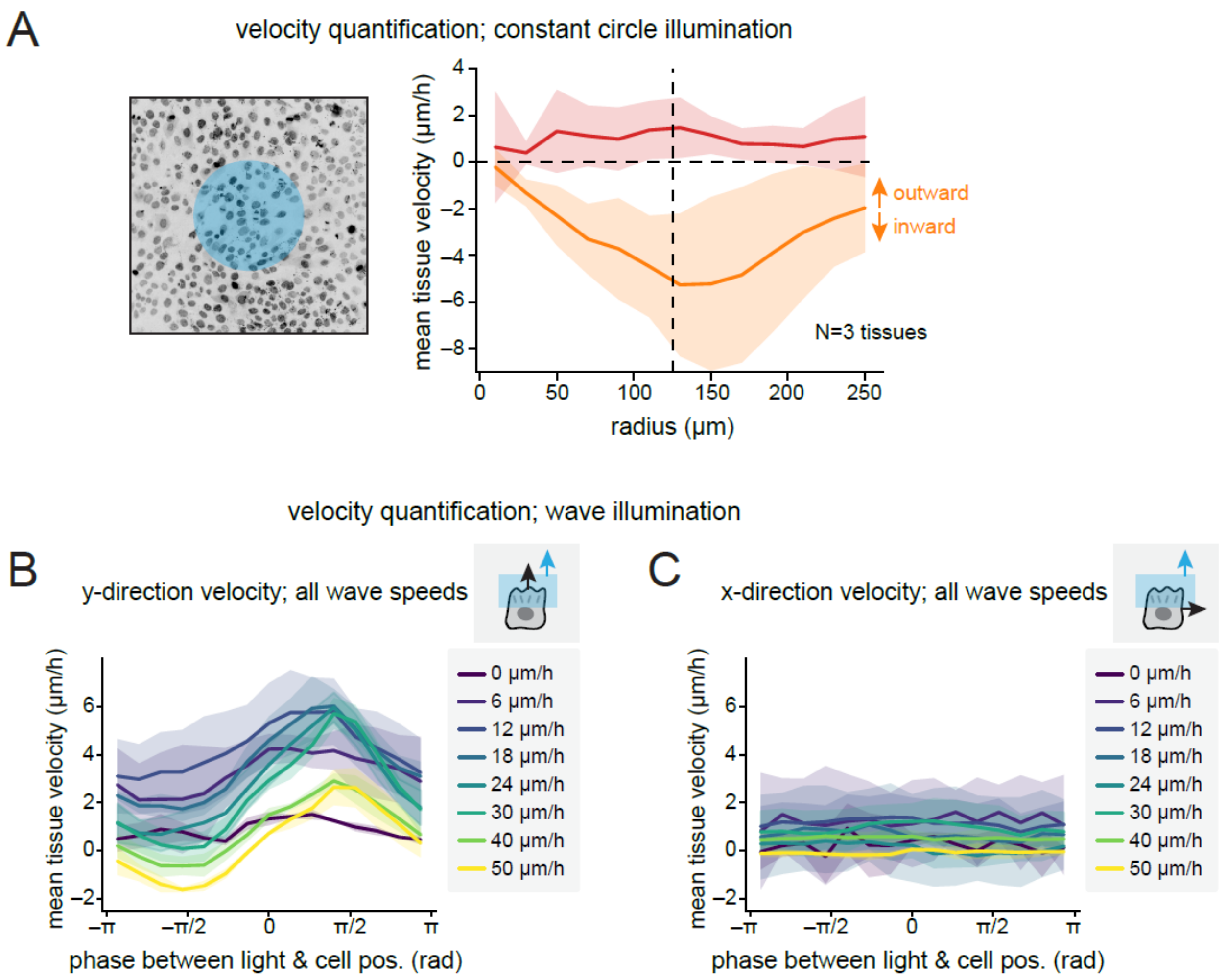


**Figure S1, related to Figure 2. Additional quantification of tissue velocities in constant-illumination and wave-illumination experiments.** (**A**) Quantification of radial and angular velocities across 3 tissue replicates. Tissues were exposed to a circle of illumination with radius 125 μm and imaged for 6 hours; data shows mean + S.D. for n=3 independent tissues. (**B-C**) Quantification of velocity in the direction of the light wave (in **B**) and perpendicular to the direction of the light wave (in **C**) as a function of light phase across all wave speeds tested. Data shows mean + S.D. across n=2,3,5,3,4,3,3,3 tissues for the 0,6,12,18,24,30,40,50 μm/h wave speeds, respectively.

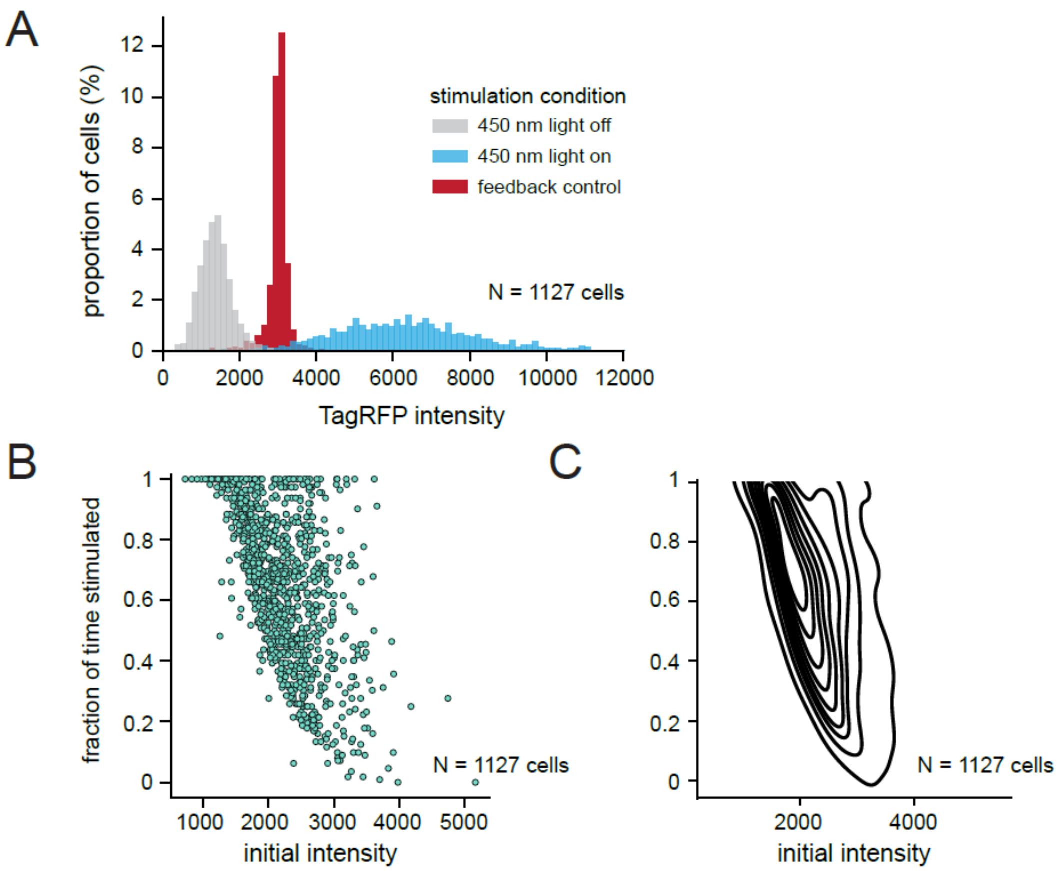


**Figure S2, related to Figure 4. Additional analyses of feedback control experiments.** (**A**) Histograms of raw TagRFP nuclear fluorescence intensity from cells before stimulation (gray), after constant 450 nm illumination (blue) and under feedback control (red). All histograms show 1127 cells from one representative experiment of n=3 replicates. (**B-C**) Quantification of the fraction of time illuminated by the feedback controller as a function of cells' initial intensities, showing results for all individual cells (in **B**) and as a contour plot (in **C**) for 1127 cells from one representative experiment of n=3 replicates.

### Supplementary Text

We provide .TOML files for all experiments. Note that any multi-point experiment can be run with multiple positions in an XML file that specifies positions whose names begin with the corresponding TOML name. For example a scheme in which three positions are chosen and named:

Position 1: circle125.1
Position 2: circle125.2
Position 3: bar12.1

will run two circle illumination patterns and one wave illumination experiment in parallel, using a single schedule.toml file to organize the overall imaging frequency and experiment duration.

See the PyCLM Github page for extensive documentation on the code and its usage, as well as updates with new stimulation experiment types.

**Circle stimulation (Fig. 2):**

schedule.toml
circle125.toml

**Traveling wave stimulation (Fig. 2):**

schedule.toml
bar12.toml
bar50.toml

(other bar speeds can be generated by editing the parameter on the final line of either TOML file)

**Vortex experiment (Fig. 3):**

schedule.toml
vortex.toml

**Global illumination cycling (Fig. 4):**

schedule.toml
vortex.toml

**Bang-bang control (Fig. 4):**

schedule.toml
clamp3000.toml

**Patterned feedback control (Fig. 4):**

schedule.toml
shield.toml

**schedule.toml**

```toml
[timing]

# number of intervals
steps = 1080

# length of the interval in seconds
interval_seconds = 60

# how far before the imaging interval to send details to the microscope (in
seconds)
setup_time_seconds = 3.0

# time in seconds between each position
# (microscope may take longer, depending on imaging time and stage movement)
time_between_positions = 3.0
```

**circle125.toml**

```
# This experiment images at 638nm and stimulates a disc in the center of the
camera

# information related to imaging conditions, channels, frequency
#  config  groups  and  device  properties  are  based  on  your  micromanager
configuration
[config_groups]
Objective = "1-Plan Apo LmbdD0.80 20x"
System = "Startup"

[device_properties]


[imaging]
every_t = 5   # image every 5 loops
exposure = 500
save = true
binning = 2

    [imaging.config_groups]
    # information related to imaging conditions, channels, frequency
    Laser-Intensity = "25%"
    LightPath = "Fluor"


# determines channel axis
[channels]
group = "Channel"
presets = ["638"]

# information related to stimulation
[stimulation]
every_t = 1   # stimulate every loop
exposure = 500
save = false

    # config groups related to stimulation acquisition
    [stimulation.config_groups]
    Channel = "DMD"
    LightPath = "DMD"
    Laser-Intensity = "AllOff"
    Sola = "30"


# information related to segmentation
[segmentation]
method = "none"


# specify the pattern method and fill in its kwargs
[pattern]
method = "circle"
rad = 125  # circle radius is 125 um
```

**bar12.toml**

```toml
# This experiment images at 638nm and moves vertical bars at a speed of 12 um/h

# information related to imaging conditions, channels, frequency
#  config  groups  and  device  properties  are  based  on  your  micromanager
configuration
[config_groups]
Objective = "1-Plan Apo LmbdD0.80 20x"
System = "Startup"

[device_properties]


[imaging]
every_t = 5   # image every 5 loops
exposure = 500
save = true
binning = 2

    [imaging.config_groups]
    # information related to imaging conditions, channels, frequency
    Laser-Intensity = "25%"
    LightPath = "Fluor"


# determines channel axis
[channels]
group = "Channel"
presets = ["638"]

# information related to stimulation
[stimulation]
every_t = 1   # stimulate every loop
exposure = 500
save = false

    # config groups related to stimulation acquisition
    [stimulation.config_groups]
    Channel = "DMD"
    LightPath = "DMD"
    Laser-Intensity = "AllOff"
    Sola = "30"


# information related to segmentation
[segmentation]
method = "none"


# specify the pattern method and fill in its kwargs
[pattern]
method = "bar"
period = 100  # spatial period: bars are placed every 100 um
duty_cycle = 0.2   # bars are 100um*0.2 = 20um tall
bar_speed = 0.2   # bars move at 12 um/h
```

**bar50.toml**

```toml
# This experiment images at 638nm and moves vertical bars at a speed of 50 um/h

# information related to imaging conditions, channels, frequency
#  config  groups  and  device  properties  are  based  on  your  micromanager
configuration
[config_groups]
Objective = "1-Plan Apo LmbdD0.80 20x"
System = "Startup"

[device_properties]


[imaging]
every_t = 5   # image every 5 loops
exposure = 500
save = true
binning = 2

    [imaging.config_groups]
    # information related to imaging conditions, channels, frequency
    Laser-Intensity = "25%"
    LightPath = "Fluor"


# determines channel axis
[channels]
group = "Channel"
presets = ["638"]

# information related to stimulation
[stimulation]
every_t = 1   # stimulate every loop
exposure = 500
save = false

    # config groups related to stimulation acquisition
    [stimulation.config_groups]
    Channel = "DMD"
    LightPath = "DMD"
    Laser-Intensity = "AllOff"
    Sola = "30"


# information related to segmentation
[segmentation]
method = "none"


# specify the pattern method and fill in its kwargs
[pattern]
method = "bar"
period = 100  # spatial period: bars are placed every 100 um
duty_cycle = 0.2  # bars are 100um*0.2 = 20um tall
bar_speed = 0.8333  # bars move at 50 um/h
```

**vortex.toml**

```toml
# This experiment images at 638nm, segments using cellpose, and
# then applies a gradient of light tangent to each cell

# information related to imaging conditions, channels, frequency
#  config  groups  and  device  properties  are  based  on  your  micromanager
configuration
[config_groups]
Objective = "1-Plan Apo LmbdD0.80 20x"
System = "Startup"

[device_properties]


[imaging]
every_t = 5  # image every 5 loops
exposure = 500
save = true
binning = 2

    [imaging.config_groups]
    # information related to imaging conditions, channels, frequency
    Laser-Intensity = "25%"
    LightPath = "Fluor"


# determines channel axis
[channels]
group = "Channel"
presets = ["638"]

# information related to stimulation
[stimulation]
every_t = 1  # stimulate every loop
exposure = 500
save = false

    # config groups related to stimulation acquisition
    [stimulation.config_groups]
    Channel = "DMD"
    LightPath = "DMD"
    Laser-Intensity = "AllOff"
    Sola = "30"


# information related to segmentation
[segmentation]
method = "cellpose"
save = true

# all additional arguments are passed to method
model = "cpsam"
gpu = true
```

```
# specify the pattern method and fill in its kwargs
[pattern]
# several per-cell stimulation options are already implemented
# this method applies asymmetric stimulation to each cell tangent to its
displacement from the center
method = "rotate_ccw"

# patterns
channel = "638"  # channel that needs to be segmented for this pattern

# options for generating the per-cell stimulation:
# voronoi determines the method for generating the per-cell stimulation region
# if voronoi is false, the stimulation is applied only on the cell masks
# (this is good for cell segmentation, bad for nuclear segmentation)
# if voronoi is true, each cell's voronoi region is used for stimulation
voronoi = true
# gradient determines the pattern of light applied in each stimulation region
# if gradient is true, a gradient of light is applied in the stimulation region
for each cell
# if gradient is false, half of the stimulation region is on, half is off
gradient = true
```

**global_cycle.toml**

```
# Part of the feedback-control experiments
# This experiment images at 545nm and 638nm, and applies global light
stimulation for 10m on, 10m off
# This experiment generates segmentations, but is open loop by design

# information related to imaging conditions, channels, frequency
[config_groups]
Objective = "1-Plan Apo LmbdD0.80 20x"
System = "Startup"

[device_properties]


# default imaging parameters, which can be overwritten by channels
[imaging]
every_t = 1   # image every loop
exposure = 200
save = true
binning = 2

    [imaging.config_groups]
    # information related to imaging conditions, channels, frequency
    Laser-Intensity = "25%"
    LightPath = "Fluor"


# determines channel axis
[channels]
group  =  "Channel"    #  config  group  that  determines  channel  (micromanager
convention)
presets = ["545", "638"]  # image in 545 and 638

    # channel specific details, overwrites imaging properties
    [channels.638]
    exposure = 100

    [channels.638.config_groups]
    Laser-Intensity = "50%"

    [channels.545]
    exposure = 200

    [channels.545.config_groups]
    Laser-Intensity = "10%"


# information related to stimulation
[stimulation]
every_t = 1
exposure = 500
save = false

    [stimulation.config_groups]
    Channel = "DMD"
```

```
    LightPath = "DMD"
    Laser-Intensity = "AllOff"
    Sola = "100"


# information related to segmentation
[segmentation]
method = "cellpose"
save = true

# all additional arguments are passed to method
model = "cpsam"
gpu = true


# specify the pattern method and fill in its kwargs
[pattern]
method = "global_cycle"

# how often to switch on-off or off-on (in minutes)
period_m = 10

# which channel to use for segmentation
nuc_channel = "545"
```

**clamp3000.toml**

```
# Part of the feedback-control experiments
# This experiment images at 545nm and 638nm, segments using cellpose, and then
stimulates
# cells below a target intensity in the 545 channel
# Closed loop experiment

# information related to imaging conditions, channels, frequency
[config_groups]
Objective = "1-Plan Apo LmbdD0.80 20x"
System = "Startup"

[device_properties]


# default imaging parameters, which can be overwritten by channels
[imaging]
every_t = 1  # image every loop
exposure = 200
save = true
binning = 2

    [imaging.config_groups]
    # information related to imaging conditions, channels, frequency
    Laser-Intensity = "25%"
    LightPath = "Fluor"


# determines channel axis
[channels]
group  =  "Channel"   #  config  group  that  determines  channel  (micromanager
convention)
presets = ["545", "638"]  # image in 545 and 638

    # channel specific details, overwrites imaging properties
    [channels.638]
    exposure = 100

    [channels.638.config_groups]
    Laser-Intensity = "50%"

    [channels.545]
    exposure = 200

    [channels.545.config_groups]
    Laser-Intensity = "10%"


# information related to stimulation
[stimulation]
every_t = 1
exposure = 500
save = false

    [stimulation.config_groups]
```

```
    Channel = "DMD"
    LightPath = "DMD"
    Laser-Intensity = "AllOff"
    Sola = "30"


# information related to segmentation
[segmentation]
method = "cellpose"
save = true

# all additional arguments are passed to method
model = "cpsam"
gpu = true


# information related to pattern generation
[pattern]
method = "binary_nucleus_clamp"

# target intensity (chosen based on imaging data from global cycle experiment)
clamp_target = 3000

# channel to segment
nuc_channel = "545"
```

**shield.toml**

```toml
# Part of the feedback-control experiments
# This experiment images at 545nm and 638nm, segments using cellpose,
# and stimulates cells below their target intensity.
# A cell's target intensity is determined by its position on a grayscale image,
placed in the center of the camera
# Closed loop experiment

# information related to imaging conditions, channels, frequency
[config_groups]
Objective = "1-Plan Apo LmbdD0.80 20x"
System = "Startup"

[device_properties]


# default imaging parameters, which can be overwritten by channels
[imaging]
every_t = 1  # image every loop
exposure = 200
save = true
binning = 2

    [imaging.config_groups]
    # information related to imaging conditions, channels, frequency
    Laser-Intensity = "25%"
    LightPath = "Fluor"


# determines channel axis
[channels]
group  =  "Channel"   #  config  group  that  determines  channel  (micromanager
convention)
presets = ["545", "638"]  # image in 545 and 638

    # channel specific details, overwrites imaging properties
    [channels.638]
    exposure = 100

    [channels.638.config_groups]
    Laser-Intensity = "50%"

    [channels.545]
    exposure = 200

    [channels.545.config_groups]
    Laser-Intensity = "10%"


# information related to stimulation
[stimulation]
every_t = 1
exposure = 500
save = false
```

```toml
    [stimulation.config_groups]
    Channel = "DMD"
    LightPath = "DMD"
    Laser-Intensity = "AllOff"
    Sola = "30"


# information related to segmentation
[segmentation]
method = "cellpose"
save = true

# all additional arguments are passed to method
model = "cpsam"
gpu = true


# information related to pattern generation
[pattern]
method = "centered_image"

# target intensities are scaled to the following values, such that the
# brightest parts of the image correspond to cells which target max_intensity,
# and the dimmest parts of the image target min_intensity
min_intensity = 2000
max_intensity = 4500

# path to target image
tif_path = "shield.tif"

# channel to segment
nuc_channel = "545"
```

**Supplementary Movie Legends**

**Movie S1.** Light-induced movement of MCF10A OptoEGFR cells toward a region of local illumination (blue circle with 125 μm radius). Left: image of cells' infrared fluorescent Hoechst nuclear dye. Right: same image with local illumination region overlaid. Scale bar 100 μm. Time shows hh:mm after stimulation. Related to Figure 2.

**Movie S2.** Light-induced movement of MCF10A OptoEGFR cells under traveling waves of illumination (bars with width 20 μm spaced every 100 μm, traveling at 12 μm/h). Left: image of cells' infrared fluorescent Hoechst nuclear dye. Right: same image with local illumination region overlaid. Scale bar 100 μm. Time shows hh:mm after stimulation. Related to Figure 2.

**Movie S3**. Light-induced movement of MCF10A OptoEGFR cells under traveling waves of illumination (bars with width 20 μm spaced every 100 μm, traveling at 50 μm/h). Left: image of cells' infrared fluorescent Hoechst nuclear dye. Right: same image with local illumination region overlaid. Scale bar 100 μm. Time shows hh:mm after stimulation. Related to Figure 2.

**Movie S4.** Light-induced movement of MCF10A OptoEGFR cells under local single-cell stimuli (blue regions in a counterclockwise orientation from the image center). Left: image of cells' infrared fluorescent Hoechst nuclear dye. Right: same image with local illumination region overlaid. Scale bar 100 μm. Time shows hh:mm after stimulation. Related to Figure 3.

**Movie S5.** Reversible stimulation of MCF10A TagRFP-NLS cells under cycles of darkness and global illumination. Left: image of cells' infrared fluorescent Hoechst nuclear dye. Right: same image with illumination region overlaid. Scale bar 100 μm. Time shows hh:mm after stimulation. Related to Figure 4.

**Movie S6.** Feedback control of MCF10A TagRFP-NLS cells. Left: image of cells' infrared fluorescent Hoechst nuclear dye. Right: same image with illumination regions overlaid. Scale bar 100 μm. Time shows hh:mm after stimulation. Related to Figure 4.

**Movie S7.** Feedback control of MCF10A TagRFP-NLS cells to a grayscale, spatially variable set point. Left: image of cells' infrared fluorescent Hoechst nuclear dye. Right: same image with illumination regions overlaid. Scale bar 100 μm. Time shows hh:mm after stimulation. Related to Figure 4.
